# A novel restraint device for injection of *Galleria mellonella* larvae that minimises the risk of accidental operator needle stick injury

**DOI:** 10.1101/085340

**Authors:** J.P. Dalton, B. Uy, S. Swift, S. Wiles

## Abstract

Larvae of the insect *Galleria mellonella* are increasingly being used for studying pathogenic microbes and their virulence mechanisms, and as a rapid model for screening novel antimicrobial agents. The larvae (waxworms) are most frequently infected by injection of pathogenic organisms into the haemocoel through the insect’s prolegs. The mostly widely used method for restraining the waxworms for injection is by grasping them between the operator’s fingers, which puts the operator at risk of needle stick injury, an important consideration when working with highly pathogenic and/or drug-resistant microorganisms. While use of a stab proof glove can reduce this risk of injury, it does so at the loss of manual dexterity and speed, resulting in a more labour-intensive and cumbersome assay. We describe a simple cost effective device (the so-called ‘*Galleria* Grabber’) for restraining waxworms for injection that keeps the operator’s fingers clear of the needle thus reducing the risk of injury.

## Introduction

Larvae (waxworms) of the Greater wax moth *Galleria melonella* have become a widely used surrogate host for studying pathogenic microbes. In recent years, they have been used for studying virulence mechanisms, investigating differences between clinical isolates as well as for preliminary investigation of the efficacy of antimicrobial compounds, for a wide range of both Gram-positive and Gram-negative bacteria^1–12^, fungi^13–19^ and viruses^20–22^. The use of waxworms as a model host has many advantages. The waxworms themselves are cheap and easy to obtain from commercial insect suppliers, and can be housed in large numbers to allow for greater study sizes at low cost. Waxworms possess an innate immune system that contains many analogous functions to that seen in humans, including phagocytosis and the production of antimicrobial peptides and reactive oxygen and nitrogen species^23^. Unlike other non-mammalian model organisms, such as *Caenorhabditis elegans, Danio rerio* and *Drosophila melanogaster*^24–21^, waxworms can be incubated at 37°C which allows for the study of clinically relevant human pathogens at a temperature that mimics the human host. Finally, as insects, *G. mellonella* are not currently subject to the same ethical restrictions that small mammalian models are, meaning there is a low barrier to entry for researchers wishing to move their studies into a model host.

Infection of waxworms is typically carried out on 5^th^ instar insects, when the waxworms are at their largest, typically around 2cm in length and 100mg in weight. The most common method of infection is by injection into the haemocoel through the last proleg of the insect; methods for injection vary between laboratories. One method is to immobilize the needle itself and then place the waxworm onto the needle for injection. Another more favoured method is to immobilise the waxworms between the operator’s fingers^28^ and place the needle into the insect’s proleg, lifting the needle away from the operator with the insect attached before pushing the plunger on the syringe. Both of these injection techniques present a hazard to the researcher and can result in needle stick injury and possible infection.

A recent article highlighted the use of a stab-proof glove to reduce the chance of this type of injury, while immobilising the waxworms over a pipette tip fixed to some paper^29^. We have tried this technique, and found that it reduced the efficiency of injection, from 3-4 infections per minute to 1 infection per minute, resulting in a lower injection rate and a more labour-intensive assay. Because of this, we investigated the possibility of using a simple restraining device to hold waxworms in place for injection, in a way that removes the operator’s hand from the vicinity of the needle, allowing for maximum mobility and safety of the operator.

## Materials and methods

### Preparation of bacteria

The *Staphylococcus aureus* isolate XEN36^30^ (Perkin Elmer) was grown overnight with shaking at 200rpm in Tryptic Soy broth (Oxoid) at 37°C. Cells were washed twice in phosphate buffered saline (PBS) (Sigma-Aldrich) and then resuspended in PBS to an optical density at 600nm (OD_600_) of 1, equivalent to approx. 5×10^9^ CFU ml^−1^. Resuspended cultures were serially diluted and plated onto Tryptic Soy agar (Oxoid) to retrospectively determine the bacterial counts used for injection. Inoculation doses were drawn into 1 ml ultra-fine (29 gauge) needle insulin syringes (BD, Wellington) for injection into the waxworms. Groups of waxworms were injected with 20 μl of either approx. 5×10^7^ CFU ml^−1^, 5×10^8^ CFU ml^−1^ or 5×10^9^ CFU ml^−1^ *S. aureus* XEN36.

### Selection, infection and monitoring of G. mellonella waxworms

5^th^ instar waxworms were selected based on consistency in size and split into eight groups of 12.Four groups were injected with either PBS or doses of 10^5^−10^7^ CFU *S. aureus* XEN36 using the most common technique of grasping the waxworms between the operator’s thumb and index finger and injecting into the waxworm’s last proleg. The remaining four groups were injected with either PBS or doses of 10^5^−10^7^ CFU *S. aureus* XEN36 using the newly described restraining device (which we have dubbed the ’*Galleria* Grabber’), which comprises a 12 cm × 9 cm kitchen sponge and a large bulldog clip (approx. 50 cm) (Fig. 1A). To comfortably restrain the waxworms, the sponge was folded in half and secured using the bulldog clip (Fig. 1B). The open ends of the folded sponge were peeled back and held in place (Fig. 1C). Next, a waxworm was placed within the sponge and held in place while the open end of the sponge was released (Fig. 1D). Once the waxworm was securely held in place, the insulin syringe was inserted into the haemocoel via the insect’s last proleg (Fig. 1E). Once the needle was in place the waxworm was released from the restraining device (Fig. 1F). If the needle is correctly placed, the waxworm remains attached to the needle of the syringe. Once the needle had been securely inserted into the waxworm, the insect was removed from the restraining device and the plunger of the syringe pushed down to inject the desired inoculum.

Once injected, waxworms were housed in individual wells of 24 well tissue culture dishes (Nunc) with the lids taped down to ensure against escape. These dishes were placed inside a secondary container to ensure containment. Waxworm mortality was monitored over 5 days.

**Figure 1.**
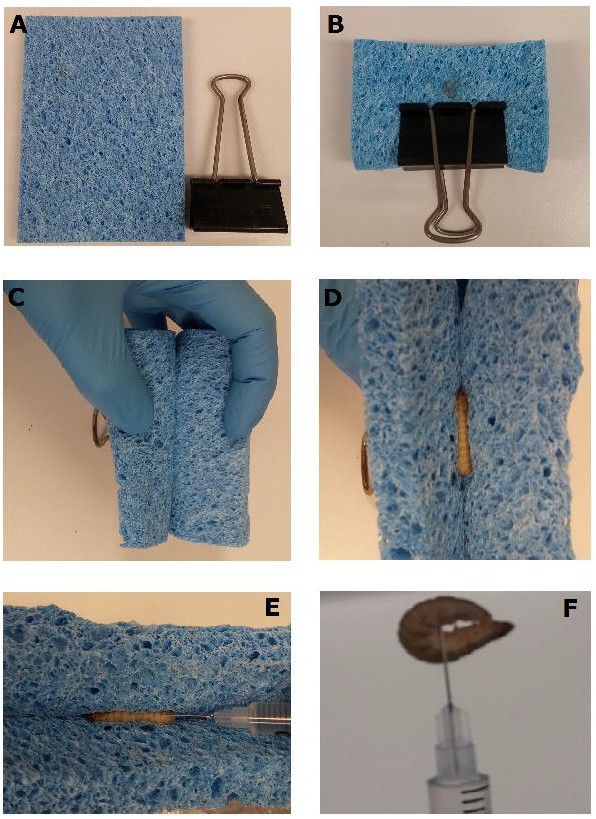
Injection of waxworms using a novel restraint device. The ‘*Galleria* Grabber’ restraint device is comprised of a 15mm thick sponge and bulldog clip (A). The sponge is folded in half lengthways and secured within a bull dog clip with the open end facing outwards (B). The open ends of the folded sponge are peeled back and held in place (C). The waxworm to be injected is placed within the sponge and held in place while the open end of the sponge is released. The closing of the sponge secures the waxworm in place for injection (E). Once the needle is placed, the syringe is lifted with the waxworm in place and the plunger is pushed to inject the desired inoculum (F).

## Results and discussion

We observed no differences in the infection dynamics between the groups of waxworms injected with *S. aureus* XEN36 after restraint using the novel ‘*Galleria* Grabber’ device described compared to restraint by holding the waxworms between the operator’s thumb and index finger. For both restraint techniques, we observed no mortality from the waxworms injected with PBS (Fig. 2). In contrast, the majority of waxworms injected with approx. 10^7^ CFU *S. aureus* XEN36 died within 24 hours (Fig. 2). We observed a dose dependent mortality for waxworms injected with *S. aureus* XEN36, with 66% of waxworms injected with approx. 10^6^ CFU succumbing to infection (Fig. 2). No mortality was seen after injection with 10^5^ CFU *S. aureus* XEN36 (Fig. 2).

The ‘*Galleria* Grabber’ allows for easy injection of a large number of waxworms (approx. 3 per minute), while greatly reducing the opportunity for the operator to suffer a needle stick injury. With the increasing popularity of waxworms as a model host for studies involving dangerous human pathogens^12^, including clinical and/or drug-resistant isolates, protecting researchers from accidental laboratory infection is of great importance. While the use of a stab-resistant glove addresses this issue, it does compromise the speed at which waxworms can be injected. With this new restraint method we were also able to inject smaller waxworms with ease. Most importantly, the new methodology described removes the operator’s hand from the vicinity of needles loaded with pathogenic/drug-resistant microbes, allowing for maximum mobility and safety of the operator without compromising the speed of the assay.

**Figure 2.**
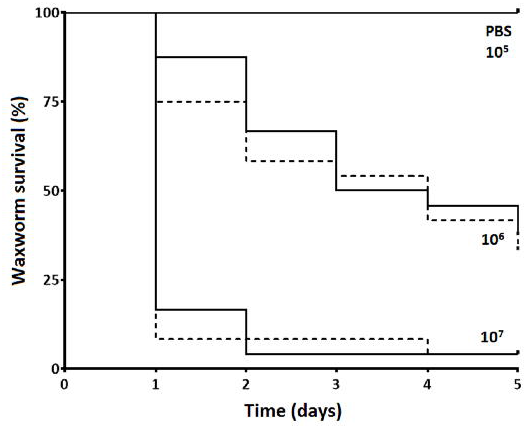
Survival of waxworms injected with varying concentrations of *S. aureus*. Waxworms (n=12 per group) were infected with varying concentrations of *S. aureus* XEN36 by injection into the haemocoel via the last proleg while restrained either between the thumb and index finger of the operator (solid lines), or using the ‘*Galleria* Grabber’ restraint device (dashed lines), and survival measured over 5 days.

## References

1 Joyce SA, Gahan CG. Molecular pathogenesis of Listeria monocytogenes in the alternative model host Galleria mellonella. Microbiology 2010; 156: 3456–3468

2 Loh JM, Adenwalla N, Wiles S etalGalleria mellonella larvae as an infection model for group A streptococcus. Virulence 2013; 4: 419–428

3 McLaughlin HP, Xiao Q, Rea RB et al A putative P-type ATPase required for virulence and resistance to haem toxicity in Listeria monocytogenes. PLOS One 2012; 7: e30928

4 Ramarao N, Nielsen-Leroux C, Lereclus D. The insect Galleria mellonella as a powerful infection model to investigate bacterial pathogenesis. J Vis Exp 2012: e4392

5 Williamson DA, Mills G, Johnson JR et al In vivo correlates of molecularly inferred virulence among extraintestinal pathogenic Escherichia coli (ExPEC) in the wax moth Galleria mellonella model system. Virulence 2014; 5: 388–393

6 Adamson DH, Krikstopaityte V, Coote PJ. Enhanced efficacy of putative efflux pump inhibitor/antibiotic combination treatments versus MDR strains of Pseudomonas aeruginosa in a Galleria mellonella in vivo infection model. J Antimicrob Chemother 2015; 70: 2271–2278

7 Thomas RJ, Hamblin KA, Armstrong SJ et al Galleria mellonella as a model system to test the pharmacokinetics and efficacy of antibiotics against Burkholderia pseudomallei. Int J Antimicrob Agents 2013; 41: 330–336

8 Moreira AS, Mil-Homens D, Sousa SA et al Variation of Burkholderia cenocepacia virulence potential during cystic fibrosis chronic lung infection. Virulence 2016: 1–15

9 Johnston T, Hendricks GL, Shen S et al Raf-kinase inhibitor GW5074 shows antibacterial activity against methicillin-resistant Staphylococcus aureus and potentiates the activity of gentamicin. Future Med Chem 2016; 8: 1941–1952

10 Nale JY, Chutia M, Carr P et al ‘Get in Early’ Biofilm and Wax Moth (Galleria mellonella) models reveal new insights into the therapeutic potential of Clostridium difficile bacteriophages. Front Microbiol 2016; 7: 1383

11 Yang H, Chen G, Hu L et al Enhanced efficacy of imipenem-colistin combination therapy against multiple-drug-resistant Enterobacter cloacae: in vitro activity and A Galleria mellonella model. J Microbiol Immunol Infect 2016, pii:S1684-1182 16 00034–00037

12 Champion OL, Wagley S, Titball RW. Galleria mellonella as a model host for microbiological and toxin research. Virulence 2016; 7: 840–845

13 De Lacorte Singulani J, Scorzoni L, De Paula E et al Evaluation of the efficacy of antifungal drugs against Paracoccidioides brasiliensis and Paracoccidioides lutzii in a Galleria mellonella model. Int J Antimicrob Agents 2016; 48: 292–297

14 Forastiero A, Bernal-Martinez L, Mellado E et al In vivo efficacy of voriconazole and posaconazole therapy in a novel invertebrate model of Aspergillus fumigatus infection. Int J Antimicrob Agents 2015; 46: 511–517

15 Frenkel M, Mandelblat M, Alastruey-Izquierdo A et al Pathogenicity of Candida albicans isolates from bloodstream and mucosal candidiasis assessed in mice and Galleria mellonella. J Mycol Med 2016; 26: 1–8

16 Borman AM, Szekely A, Johnson EM. Comparative pathogenicity of United Kingdom isolates of the emerging pathogen Candida auris and other key pathogenic Candida species. Msphere 2016; 14. pii:e00189–e00e16

17 Gago S, Serrano C, Alastruey-Izquierdo A et al Molecular identification, antifungal resistance and virulence of Cryptococcus neoformans and Cryptococcus deneoformans isolated in Seville, Spain. Mycoses 2016 doi:10.1111/myc.12543. [Epub ahead of print].

18 Santos R, Costa C, Mil-Homens D et al The multidrug resistance transporters CgTpo1_1 and CgTpo1_2 play a role in virulence and biofilm formation in the human pathogen Candida glabrata. Cell Microbiol 2016 doi:10.1111/cmi.12686. [Epub ahead of print].

19 Alcazar-Fuoli L, Buitrago M, Gomez-Lopez A et al An alternative host model of a mixed fungal infection by azole susceptible and resistant Aspergillus spp strains. Virulence 2015; 6: 376–384

20 Buyukguzel E, Tunaz H, Stanley D et al Eicosanoids mediate Galleria mellonella cellular immune response to viral infection. J Insect Physiol 2007; 53: 99–105

21 Ozkan S, Coutts RH. Aspergillus fumigatus mycovirus causes mild hypervirulent effect on pathogenicity when tested on Galleria mellonella. Fungal Genet Biol 2015; 76: 20–26

22 Garzon S, Charpentier G, Kurstak E Morphogenesis of the nodamura virus in the larbae of the lepidopteran Galleria mellonella (L.). Arch Virol 1978; 56: 61–76

23 Wojda I. Immunity of the greater wax moth Galleria mellonella. Insect Sci 2016, doi:10.1111/1744-7917.12325. [Epub ahead of print].

24 Lopez Hernandez Y, Yero D, Pinos-Rodriguez JM et al Animals devoid of pulmonary system as infection models in the study of lung bacterial pathogens. Front Microbiol 2015; 6: 38

25 Panayidou S, Ioannidou E, Apidianakis Y. Human pathogenic bacteria, fungi, and viruses in Drosophila: disease modeling, lessons, and shortcomings. Virulence 2014; 5: 253–269

26 Arvanitis M, Glavis-Bloom J, Mylonakis E. Invertebrate models of fungal infection. Biochim Biophys Acta 2013; 1832: 1378–1383

27 Glavis-Bloom J, Muhammed M, Mylonakis E. Of model hosts and man: using Caenorhabditis elegans, Drosophila melanogaster and Galleria mellonella as model hosts for infectious disease research. Adv Exp Med Biol 2012; 710: 11–17

28 Fuchs BB, O’Brien E, El Khoury JB et al Methods for using Galleria mellonella as a model host to study fungal pathogenesis. Virulence 2010; 1: 475–482

29 Harding CR, Schroeder GN, Collins JW et al Use of Galleria mellonella as a model organism to study Legionella pneumophila infection. J Vis Exp 2013: e50964

30 Francis KP, Joh D, Bellinger-Kawahara C et al Monitoring bioluminescent Staphylococcus aureus infections in living mice using a novel luxABCDE construct. Infect Immun 2000; 68: 3594–3600

